# The effect of constraining mediolateral ankle moments and foot placement on the use of the counter-rotation mechanism during walking

**DOI:** 10.1101/2021.12.14.472551

**Authors:** Maud van den Bogaart, Sjoerd M. Bruijn, Joke Spildooren, Jaap H. van Dieën, Pieter Meyns

**Affiliations:** Rehabilitation Research Center (REVAL), Faculty of Rehabilitation Sciences, Hasselt University, Diepenbeek 3590, Belgium; Department of Human Movement Sciences, Vrije Universiteit Amsterdam, Amsterdam Movement Sciences, Amsterdam, Netherlands; Institute of Brain and Behavior Amsterdam, Amsterdam, The Netherlands

**Keywords:** Gait stability, constrained walking, center of mass acceleration, foot placement, ankle moments, counter-rotation mechanism

## Abstract

Stability during walking can be maintained by shifts of the Center of Pressure through modulation of foot placement and ankle moments (CoP-mechanism). An additional mechanism to stabilize gait, is the counter-rotation mechanism i.e. changing the angular momentum of segments around the Center of Mass (CoM) to change the direction of the ground reaction force. It is unknown if and how humans use the counter-rotation mechanism to control the CoM during walking and how this interacts with the CoP-mechanism. Thirteen healthy adults walked on a treadmill, while full-body kinematic and force plate data were obtained. The contributions of the CoP and the counter-rotation mechanisms to control the CoM were calculated during steady-state walking, walking on LesSchuh, i.e. constraining mediolateral CoP shifts underneath the stance foot and walking on LesSchuh at 50% of normal step width, constraining both foot placement and ankle mechanisms (LesSchuh50%). A decreased magnitude of within-stride control by the CoP-mechanism was compensated for by an increased magnitude of within-stride control by the counter-rotation mechanism during LesSchuh50% compared to steady-state walking. This suggests that the counter-rotation mechanism is used to stabilize gait when needed. However, the mean contribution of the counter-rotation mechanism over strides did not increase during LesSchuh50% compared to steady-state walking. The CoP-mechanism was the main contributor to the total CoM acceleration. The use of the counter-rotation mechanism may be limited because angular accelerations ultimately need to be reversed and because of interference with other task constraints, such as head stabilization and preventing interference with the gait pattern.

## Introduction

Stable gait, defined as gait that does not lead to falls(Bruijn et al., 2013), requires control of the state of the body center of mass (CoM) relative to the base of support (BoS), i.e. the area within an outline of all points on the body in contact with the support surface, or vice versa. In gait, the BoS is formed by the parts of the feet that are in contact with the floor at any point in time(Bruijn and van Dieen, 2018). The most extensively studied mechanism to stabilize gait is foot placement(Bauby and Kuo, 2000, Wang and Srinivasan, 2014, Rankin et al., 2014, Bruijn and van Dieen, 2018). A second mechanism to stabilize gait is the application of moments around the ankle of the stance foot (i.e. ankle mechanism)(Horak and Nashner, 1986). These ankle moments are reflected in a shift of the center of pressure of the ground reaction force (CoP). Foot placement primarily moves the BoS to accommodate the state of the CoM, but also constrains the location of the CoP. So, foot placement and ankle moments combined determine the resulting position of the CoP relative to the CoM, which in turn determines the acceleration of the CoM and thus allows control of the CoM relative to the BoS. Shifts of the CoP through modulation of foot placement and ankle moments will here be referred to as the CoP-mechanism. An additional mechanism to stabilize gait, is the counter-rotation mechanism, i.e. changing the angular momentum of segments around the CoM to change the direction of the ground reaction force(Hof, 2007, Otten, 1999, Hof et al., 2007). A leftward rotational acceleration of the trunk, for example, results in a rightward acceleration of the CoM and vice versa. Accelerations of other body segments, for example the arms or head can be used in the same way.

The main mechanism for active control in the frontal plane has been suggested to be foot placement, with a smaller role for the ankle mechanism(Bauby and Kuo, 2000, MacKinnon and Winter, 1993, van Leeuwen et al., 2021). Theoretically, the counter-rotation mechanism may also play a role, but research on the role of the counter-rotation mechanism is limited. In a previous study, we investigated the relative contribution of the CoP-mechanism and the counter-rotation mechanism to control the CoM in the anteroposterior direction during a normal step and during the first recovery step after a perturbation (belt acceleration) in healthy adults(van den Bogaart et al., 2020). A limited use of the counter-rotation mechanism to control the CoM was found, as using the counter-rotation mechanism would actually have interfered with the gait pattern(van den Bogaart et al., 2020). The current study will address the role of the counter-rotation mechanism for mediolateral stability.

When aiming to improve gait stability, it is important to unravel the use of the CoP and counter-rotation mechanisms and their interplay. Simply studying unconstrained walking may not be sufficient, as during normal walking, the foot placement mechanism is the dominant mechanism(Bauby and Kuo, 2000, MacKinnon and Winter, 1993, van Leeuwen et al., 2021). Constrained or perturbed walking might provide insight in the use of the CoP and counter-rotation mechanisms and their interplay. Previous research has investigated the interplay between the two components of the CoP-mechanism, i.e. shifts of the CoP through modulation of foot placement and ankle moments, by constraining one of them. Ankle moment control during single stance did not compensate for constrained foot placement to a fixed step width(van Leeuwen et al., 2021). Therefore, it is likely that the counter-rotation mechanism played a role in controlling the CoM relative to the BoS. This was, however, not investigated. Conversely, constraining ankle moments, e.g. by narrowing the surface area underneath the shoe or by walking on prosthetic legs, led to a compensatory wider foot placement(Hof et al., 2007, van Leeuwen et al., 2020). This is consistent with modeling results, suggesting that without any CoP modulation during the stance phase following a sensory perturbation, a 60% wider foot placement is needed to maintain stable in the mediolateral direction(Reimann et al., 2017). Nevertheless, walking with constrained ankle moments due to a prosthetic leg did not lead to a significant increase in the range of the frontal plane angular momentum (<15%)(Silverman and Neptune, 2011, D’Andrea et al., 2014, Miller et al., 2018, Pickle et al., 2014, Sheehan et al., 2015). Assuming similar timing of the minima and maxima of the angular momentum, increasing (decreasing) the angular momentum corresponds to increased (decreased) rate of change of the angular momentum, and therefore suggests increased (decreased) use of the counter-rotation mechanism. Thus, most likely, the counter-rotation mechanism did not, or only minimally accommodate for limited ankle moments in people with a prosthetic leg. These previous studies determined mean range of the angular momentum over all strides. However, the magnitude of within-stride control by the counter-rotation mechanism and stride-to-stride variation in the contribution of the counter-rotation mechanism could still be increased to compensate for reduced modulation of ankle moments, but this was not reported. Constraints in the CoP-mechanism could be adjusted by such an increase in variation. Therefore, we will determine the magnitude of the within-stride control by the counter-rotation mechanism and stride-to-stride variation in contribution of the counter-rotation mechanism in the current study. Additionally, to study if and how humans use a counter-rotation mechanism to control the CoM during walking, it may be needed to provoke them to do so, for example by constraining foot placement in addition to constraining ankle moments. Studies that assessed the use of the counter-rotation mechanism after medial foot placement or platform perturbations when ankle moments were constrained, did find increased whole-body angular momentum(Miller et al., 2018, Sheehan et al., 2015). It seemed that by imposing restrictions on the usability of both components of the CoP-mechanism, the use of the counter-rotation mechanism could be provoked(Miller et al., 2018). The authors interpreted this increased whole-body angular momentum, however, as a sign of instability(Miller et al., 2018, Sheehan et al., 2015). However, it could be that the increased angular momentum is a sign of increased control of the CoM through the counter-rotation mechanism.

In the current study, we aim to assess the contribution of the CoP and counter-rotation mechanisms in mediolateral direction during steady-state walking and during walking with constrained ankle moments or during walking with constrained ankle moments combined with constrained foot placement (i.e. constrained CoP-mechanism). We expected that the mean contribution of the counter-rotation mechanism over strides would be similar during all conditions, because angular accelerations ultimately need to be reversed, leading to the opposite effect on the acceleration of the body center of mass. We also expected limited use of the counter-rotation mechanism compared to the CoP-mechanism, because rotational accelerations of body parts may interfere with other task constraints, such as stabilizing the orientation of the head in space and performing leg swing(van den Bogaart et al., 2020). Moreover, we expected that the within-stride control by the counter-rotation mechanism and stride-to-stride variation will increase when ankle moments are constrained, and even more when also foot placement is constrained. This hypothesis is based on a previous study, showing an increased range of whole-body angular momentum when stable gait is compromised in the presence of mediolateral foot placement perturbations in people with a prosthetic leg(Miller et al., 2018).

## Methods

### Subjects

Thirteen healthy adults (6 males, age 23.8 ± 3.7 years old, BMI 23.3 ± 2.2 kg/m^2^) participated in this study. Sample size was calculated for a two-tailed paired sample t-test analysis using G*Power (1-β = 0.9, α = 0.05) and an effect size of 1.13(Miller et al., 2018).

Potential participants were excluded if they reported any neurological or orthopedic disorder(s), had an uncorrectable visual impairment, were unable to walk without difficulty for ≥45 minutes, had undergone surgery of the lower extremities during the last two years, or took medication that might affect the gait pattern. Participants gave written informed consent prior to the experiment. Ethical approval (VCWE-2020-202) was granted by the ethics review board of the faculty of Behavioral and Movement Sciences at ‘Vrije Universiteit Amsterdam’.

### Materials and software

Participants walked on an instrumented dual-belt treadmill (Motek-Forcelink, Amsterdam, Netherlands). Full body kinematics were measured using two Optotrak cameras (Northern Digital Inc, Waterloo Ontario, Canada) directed at the center of the treadmill (sampling rate was 50 Hz). This system was calibrated with the +x axis defined to the right, the +y axis defined as forward, and the +z axis defined as upwards. Clusters of 3 markers were attached to the shoes (feet), shanks, thighs, pelvis, trunk, upper arms, forearms and the head. Corresponding anatomical landmarks were digitized using a four-marker probe.

During the experiment participants wore shoes to constrain the mediolateral shift of the CoP underneath the stance foot through a narrow (1 cm) ridge attached to the sole(Figure 1). The so-called LesSchuh limits ankle moments, while anteroposterior roll-off and subsequent push-off remain possible, because the material of the ridge bends with the sole in anteroposterior direction.

**Figure 1.**
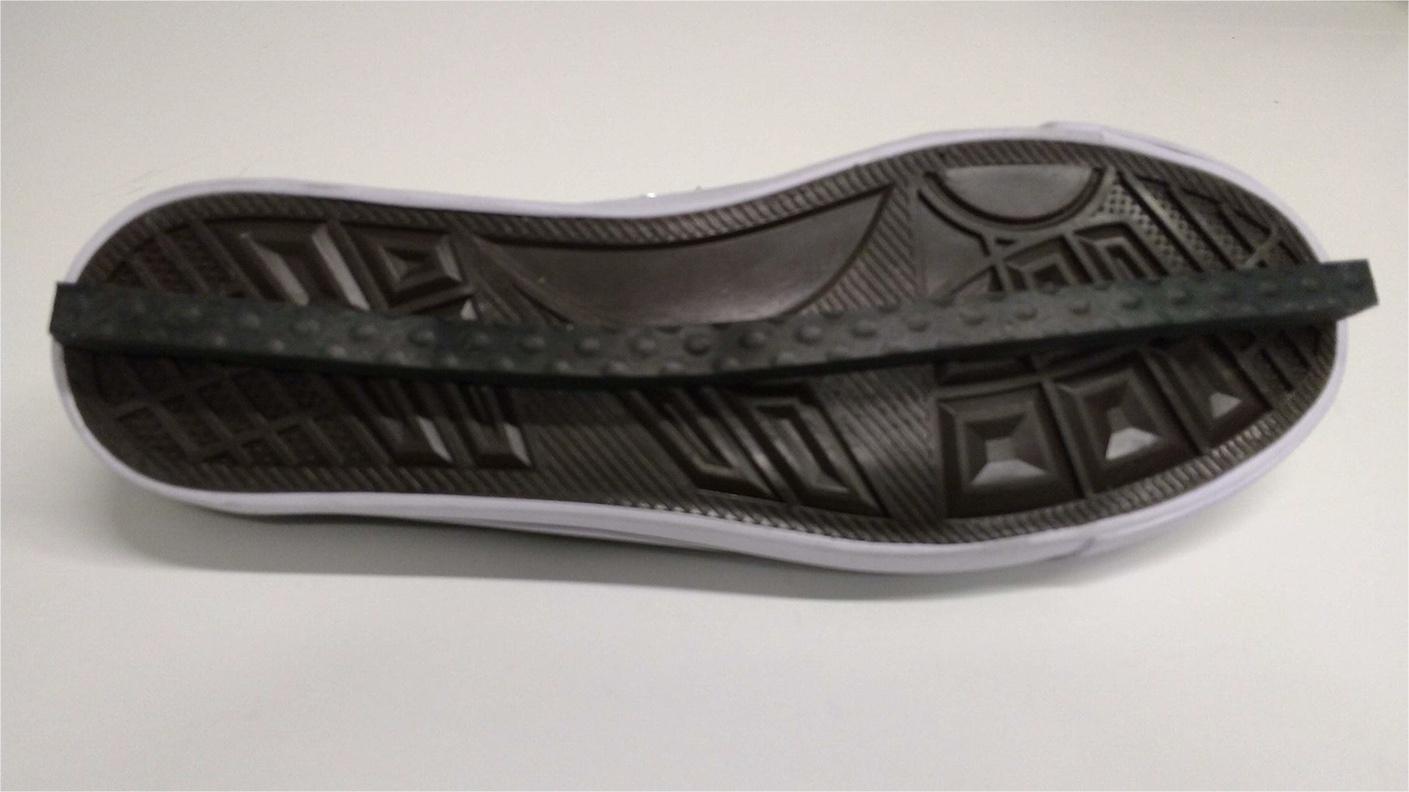
LesSchuh, shoe constraining the mediolateral shift of the CoP underneath the stance foot through the narrow (1cm) ridge attached to the sole.

### Procedures

Participants walked on the left belt of an instrumented dual-belt treadmill during three conditions at a constant speed of 2.5 km/h. Participants were required to walk on one belt to prevent that the LesSchuh could get stuck in the gap between the two belts. Additionally, walking on two belts could result in a wider step as a precaution(van Leeuwen et al., 2020). The first condition consisted of 10 minutes steady-state walking on normal shoes of the same type as the LesSchuh but without the ridge. The second condition consisted of 15 minutes walking on the LesSchuh(Figure 1). Participants were asked to walk on the ridge, not touching the ground with the sides of the shoe’s sole. Participants were also instructed to place their feet in a similar orientation as they would do without LesSchuh, to avoid a “toeing-out strategy” potentially inducing a mediolateral shift of the center of pressure after foot placement (heel strike) despite the narrow base of support(Rebula et al., 2017). The third and final condition consisted of 5 minutes walking on the LesSchuh at 50% of the average step width as measured during the first condition, constraining both components of the CoP-mechanism (i.e., foot placement and ankle mechanisms)(LesSchuh50%). The step width during the third condition was imposed by projecting beams on the treadmill, and participants were instructed to place their foot (ridge of the Lesschuh) in the middle of the beam. During all conditions, participants wore a safety harness connected to a rail at the ceiling, that did not provide weight support.

### Data analysis

Kinematic and force plate data were low-pass filtered at 10 Hz using a second order Butterworth filter. Kinematic data were analyzed using a 13-segment kinematic model. Full-body CoM was calculated by combining the CoM of all segments. CoP and full-body CoM data were high-pass filtered at 0.1 Hz using a second order Butterworth filter. Heel-strikes and toe-offs were determined based on CoP data(Roerdink et al., 2008). One full gait cycle started at heel-strike of the left foot and ended at heel-strike of the same foot (left).

The contributions of the CoP-mechanism and counter-rotation mechanism to the CoM acceleration in the frontal plane, were calculated using Eq.(1), as described by Hof (2007).

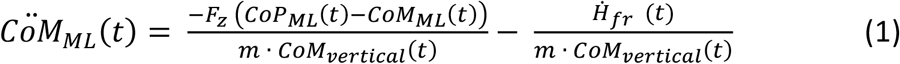

in which *m* is body mass, *CoM*_*ML*_ is the mediolateral (ML) position of the CoM, *CoM*_*vertical*_ is the vertical position of the CoM, *CöM*_*ML*_ is the double derivative of CoM_ML_ with respect to time, *t* is time, *F*_*z*_ is the vertical ground reaction force, *CoP*_*ML*_ is the ML position of the CoP, and *Ḣ*_*fr*_ is the change in total body angular momentum (around the CoM) in the frontal plane. Here the first part of the right-hand term, 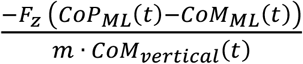, refers to the ML CoM acceleration induced by the CoP-mechanism, whereas the second part, 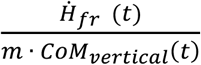, is the ML CoM acceleration induced by the counter-rotation mechanism.

For each individual stride, the time normalized curves of the total CoM acceleration and of the contributions of the CoP and counter-rotation mechanisms to the total CoM acceleration in the frontal plane were calculated. For each of these variables, we calculated the averaged curve across all strides during each trial, to obtain the average time normalized curve during a stride per trial.

We determined the magnitude of within-stride control for each of the three variables by calculating the standard deviation (SD) of the time series for each single stance phase. Then we averaged across all single stance phases during each trial, to obtain the average variability within single stance per trial. The magnitude of within-stride control by the CoP-mechanism can only be explained by the ankle mechanism, as the magnitude of within-stride control was determined during single stance. The magnitude of within-stride control of the total CoM acceleration is an index of the total amount of control of the CoM during single stance.

We also determined the magnitude of between-stride control (i.e. stride-to-stride variation) for each of the three variables by calculating the SD over the strides per time sample, after which the average SD was calculated per trial.

To determine to role of foot placement to the contribution of the CoP-mechanism, we assessed the step width per step as the difference between the mediolateral CoP positions halfway during two subsequent single leg stances and calculated the average step width across all steps per trial.

### Statistics

Average time normalized curves of the total CoM acceleration and of CoM acceleration induced by the CoP and counter-rotation mechanism were compared between conditions using SPM repeated measures ANOVA (SPM1d vM.0.4.5, www.spm1D.org). Parametric statistical tests were used, as the D’Agostino-Pearson K2 test revealed that the values were normally distributed. If the main effect was significant, post-hoc SPM (t) maps were calculated. A Bonferroni correction was applied for comparisons within one variable.

One-way repeated measures ANOVAs were used to determine the effect of Condition on the magnitude of within- and between-stride control of the total CoM acceleration and of the CoM accelerations due to the CoP and counter-rotation mechanism and on the average step width. The data were normally distributed as tested with the Shapiro-Wilk test. Post-hoc analyses were performed to determine differences between the conditions (using a Bonferroni correction). Except the SPM analyses, statistical analyses were performed with SPSS (v25), and for all analysis we used α<0.05.

## Results

### Total CoM acceleration

The average time normalized curve of the total CoM acceleration was similar during steady-state walking and walking on LesSchuh, but was decreased around left and right heel-strike during LesSchuh50% compared to steady-state walking and was increased during the single stance phases during LesSchuh compared to LesSchuh50%(Figure 2). The magnitude of within-stride control of the total CoM acceleration did not differ significantly between the conditions (*F*(2,24) = 2.494, *p* = 0.104)(Figure 3A). The magnitude of between-stride control of the total CoM acceleration differed significantly between the conditions (*F*(2,24) = 43.562, *p* < 0.001). The magnitude of between-stride control was increased during LesSchuh and LesSchuh50% compared to steady-state walking (p < 0.001 and *p* < 0.001, respectively)(Figure 3B).

**Figure 2.**
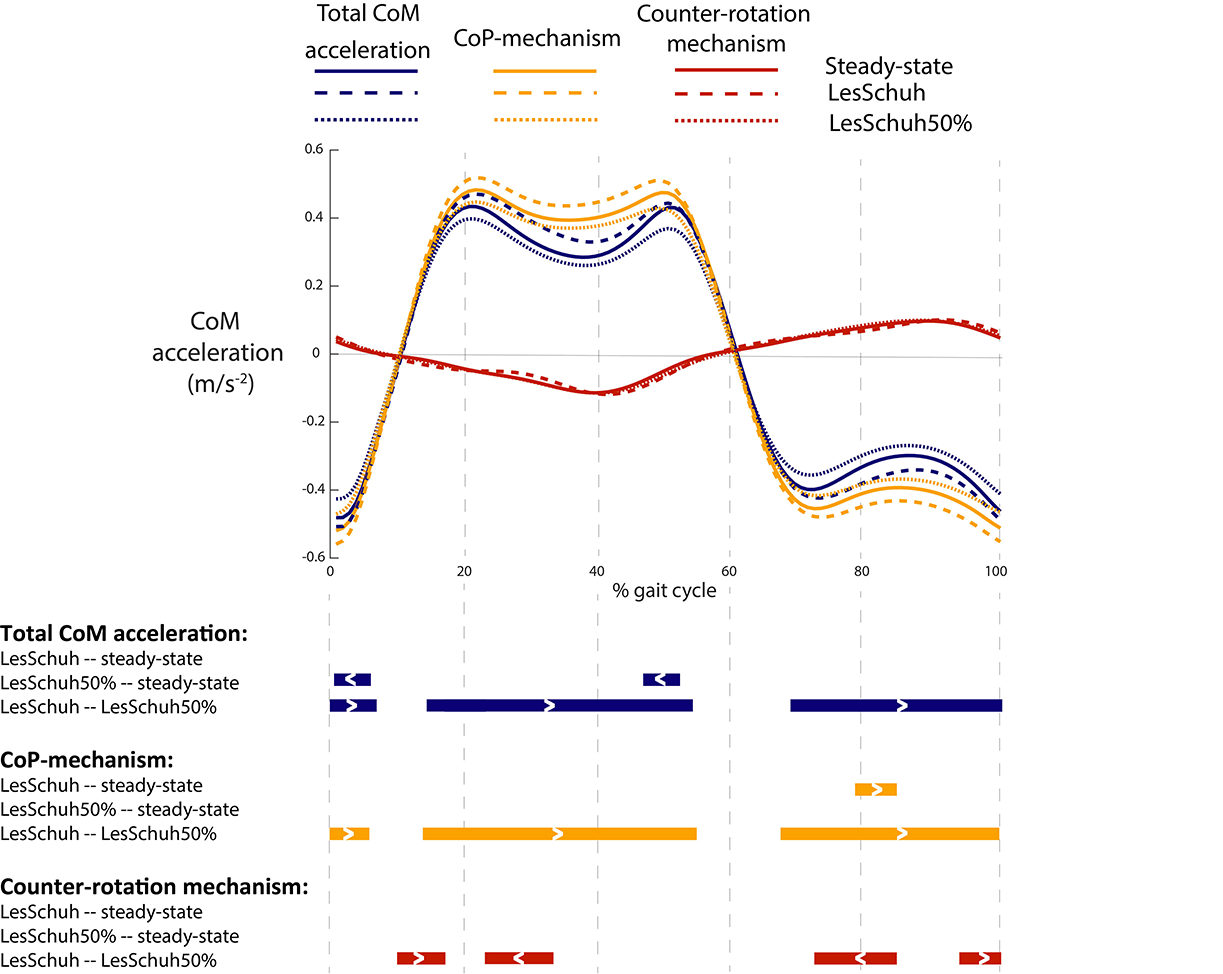
Average time series (N=13) of the center of Mass (CoM) acceleration (in blue) and the contribution of the center of pressure mechanism (CoP-mechanism) (in yellow) and the counter-rotation mechanism (in red) to the CoM acceleration during normal walking (steady-state, solid lines), walking on LesSchuh constraining the ankle mechanism (LesSchuh, dashed lines) and walking on LesSchuh at 50% of the average step width during steady-state walking constraining the ankle mechanism and foot placement (LesSchuh50%, dotted lines). The gait cycle (0-100%) started at heel-strike of the left foot (0%) and ended at heel-strike of the same foot (left) (100%). The bars indicate gait phases with significant differences between conditions. The greater then (>) and less then (<) sign within the bars indicate if the CoM acceleration during the first mentioned condition is greater or less then during the latter condition.

**Figure 3.**
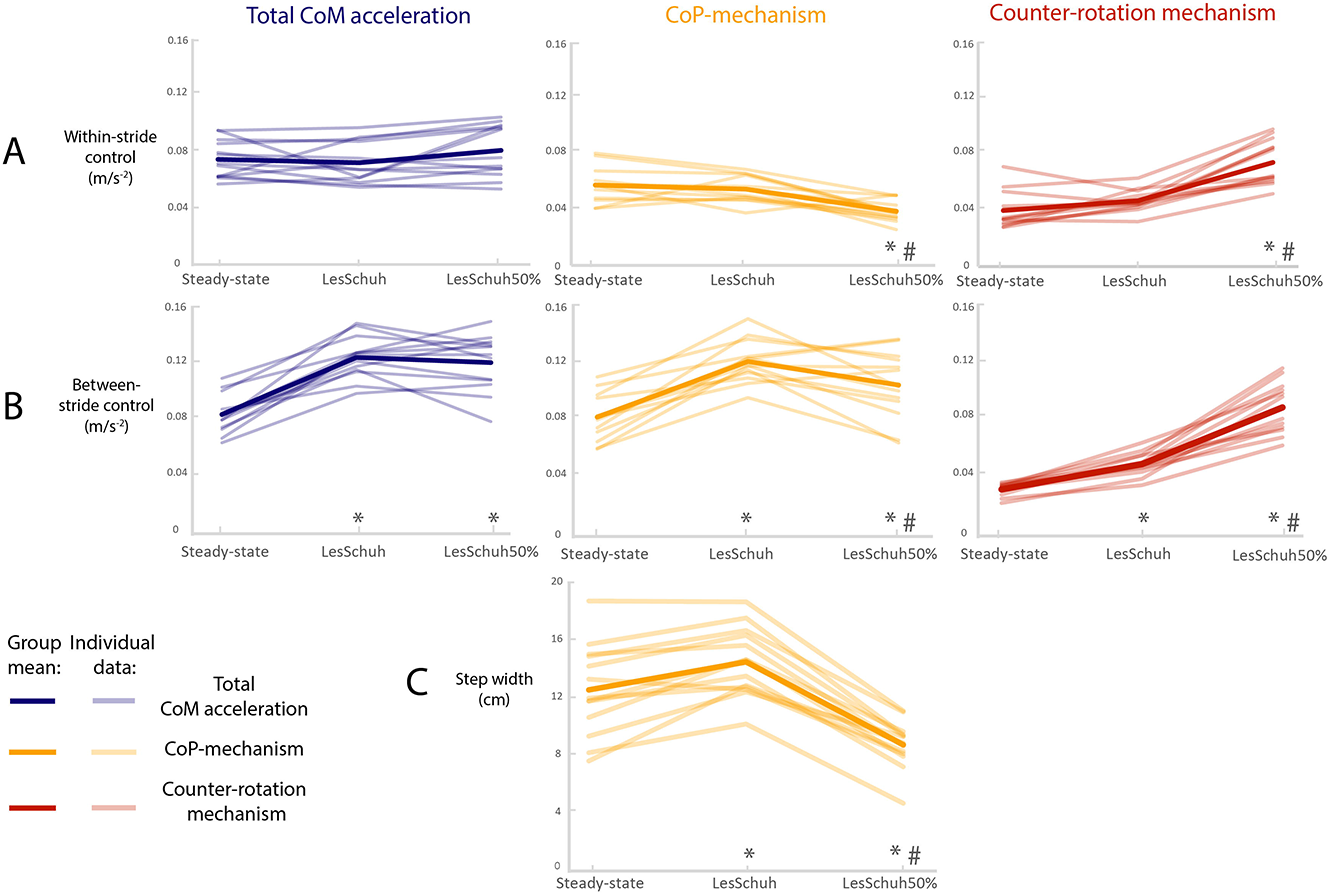
The effect of Condition on A) the magnitude of within-stride control during single stance, as reflected in the total CoM acceleration (in blue) and CoM acceleration due to the CoP-mechanism (in yellow) and due to the counter-rotation mechanism (in red), B) the magnitude of between-stride control, as reflected in the total CoM acceleration and CoM acceleration due to the CoP-mechanism and due to the counter-rotation mechanism and C) the average step width. * represents a significant difference compared to steady-state walking. # represents significant a difference compared to LesSchuh.

### CoP-mechanism

The average time normalized curve of the contribution of the CoP-mechanism to the total CoM acceleration, was increased around 80% of the gait cycle during LesSchuh compared to steady-state walking, was similar during LesSchuh50% and steady-state walking and was increased during the single stance phases during LesSchuh compared to LesSchuh50%(Figure 2).

The magnitude of within-stride control by the CoP-mechanism during single stance due to the ankle mechanism differed significantly between the conditions (*F*(2, 24) = 20.411, *p* < 0.001). The magnitude of within-stride control by the CoP-mechanism was decreased during LesSchuh50% compared to steady-state walking and LesSchuh (*p* < 0.001 and *p* = 0.001, respectively)(Figure 3A). The magnitude of between-stride control by the CoP-mechanism differed significantly between the conditions (*F*(2, 24) = 24.811, *p* < 0.001).

The magnitude of between-stride control by the CoP-mechanism was increased during LesSchuh50% and LesSchuh compared to steady-state walking (*p* = 0.011 and *p* < 0.001, respectively) and was decreased during LesSchuh50% compared to LesSchuh (*p* = 0.045)(Figure 3B). Step width differed significantly between the conditions (*F*(1.316, 15.793) = 58.234, *p* < 0.001).

Step width was decreased during LesSchuh50% compared to LesSchuh and steady-state walking (with around 28%)(*p* < 0.001 and *p* = 0.001, respectively)(Figure 3C). Step width was increased during LesSchuh compared to steady-state (*p* = 0.003)(Figure 3C).

### Counter-rotation mechanism

The average time normalized curve of the contribution of the counter-rotation mechanism to the total CoM acceleration was similar when comparing steady-state walking with LesSchuh and LesSchuh50%(Figure 2). The contribution of the counter-rotation mechanism to the total CoM acceleration was increased around 15% and 95% of the gait cycle, and decreased around 30% and 80% of the gait cycle during LesSchuh compared to LesSchuh50%(Figure 2). The magnitude of within-stride control by the counter-rotation mechanism differed significantly between the conditions (*F*(2,24) = 46.148, *p* < 0.001). The magnitude of within-stride control by the counter-rotation mechanism was increased during LesSchuh50% compared to steady-state walking and LesSchuh (*p* < 0.001 and *p* < 0.001, respectively)(Figure 3A). The magnitude of between-stride control by the counter-rotation mechanism differed significantly between the conditions (*F*(1.210, 14.515) = 120.775, *p* < 0.001). The magnitude of between-stride control by the counter-rotation mechanism increased from steady-state walking to LesSchuh to LesSchuh50% (*p* < 0.001, *p* < 0.001, *p* < 0.001, respectively)(Figure 3B).

## Discussion

We assessed the contribution of both mechanisms in normal walking and in conditions in which ankle moments (using a customized shoe (LesSchuh)) and foot placement were constrained in the mediolateral direction.

Walking on LesSchuh led to an increased step width compared to steady-state walking. This is consistent with previous literature, showing that foot placement control can accommodate for limited ankle moments in case of, e.g. narrowing the surface area underneath the shoe or prosthetic legs(Hof et al., 2007, van Leeuwen et al., 2020). We expected that the magnitude of within-stride control by the CoP-mechanism, for which only the ankle mechanism can be responsible, would decrease during both conditions wearing LesSchuh compared to steady-state walking. However, we found that the magnitude of within-stride control by the CoP-mechanism only significantly decreased during walking with ankle and foot placement constraints (LesSchuh50%) compared to steady-state walking. Toeing-out walking could be a possible explanation for the fact that it was possible to shift the CoP over a larger area in mediolateral direction. Despite the instruction to place the feet facing forward, especially one participant walked with a “toeing-out strategy” during LesSchuh. There was still no significant difference in within-stride control by the CoP-mechanism between LesSchuh and steady-state walking when running the statistical analysis without this participant. This suggests that more participants probably walked with a toeing-out strategy, which was not directly visible when visually inspecting the data.

Moreover, the magnitude of within-stride control by the counter-rotation mechanism was similar during walking on LesSchuh and steady-state walking. It seems that only constraining the ankle moments was not sufficient to provoke an increased contribution of the counter-rotation mechanism compared to steady-state walking. It appears not to be necessary to use the counter-rotation mechanism in addition to adjusting foot placement control (i.e. walking with wider steps) to control stability in these conditions. As expected, we found decreased step width and decreased within-stride control by the ankle mechanism, during LesSchuh50% compared to steady-state walking and LesSchuh. The magnitude of the within-stride and between-stride control by the counter-rotation mechanism increased during LesSchuh50% compared to steady-state walking and LesSchuh, which is consistent with our hypothesis. Compared to steady-state walking and LesSchuh, during LesSchuh50% the decreased magnitude of within-stride control by the CoP-mechanism was thus compensated for by increased magnitude of within-stride control by the counter-rotation mechanism. This compensation resulted in no difference in within-stride control of the total CoM acceleration, which is an index of the total amount of control of the CoM, between the conditions.

The increased within-stride control by the counter-rotation mechanism did not coincide with an increased mean contribution of the counter-rotation mechanism to CoM acceleration over strides during LesSchuh50% compared to steady-state walking and LesSchuh, and even a decrease around 15% and 95% of the gait cycle during LesSchuh50% compared to LesSchuh. The absence of significant differences between the conditions in mean contribution of the counter-rotation mechanism to total CoM acceleration over strides may be because angular accelerations, with the possible exception of angular acceleration of the arm around the shoulder, ultimately need to be reversed, leading to the opposite effect on the acceleration of the body center of mass and thus allowing relatively high-frequency (within stride) modulation only. Overall, the mean contribution of the counter-rotation mechanism to CoM acceleration over strides was around three times smaller and in opposite direction compared to the mean contribution of the CoP-mechanism over strides, while the magnitudes of the within-stride control by the CoP and counter-rotation mechanisms were more or less similar. This is consistent with previous findings, where we also found around three times smaller and counteracting contributions of the counter-rotation mechanism and CoP-mechanisms to control the CoM in the anteroposterior direction during a normal step and the first recovery step after perturbation(van den Bogaart et al., 2020). The CoP-mechanism was the main contributor to the total CoM acceleration. The use of the counter-rotation mechanism may interfere with other task constraints, such as stabilizing the orientation of the head in space and preventing interference with the gait pattern.

In future studies, it is worthwhile to assess the use of the counter-rotation mechanism in children, older adults or people with varying pathologies. These populations might use the balance control mechanisms differently to stabilize mediolateral or anteroposterior gait. Future studies should determine the link between falling and the use of the counter-rotation mechanism. Training the use of specific mechanisms to control the CoM could be implemented in therapeutic interventions that aim to decrease fall incidence. However, whether and how a specific mechanism can be trained needs further investigation.

## Conclusion

We found that a decreased magnitude of within-stride control by the CoP-mechanism was compensated for by an increased magnitude of within-stride control by the counter-rotation mechanism during LesSchuh50% compared to steady-state walking. This suggests that the counter-rotation mechanism is used to stabilize gait when needed. In real life, such situations may for example involve near balance loss, where a further shift of the CoP under the stance foot is impossible. However, the mean contribution of the counter-rotation mechanism to CoM acceleration over strides did not increase during LesSchuh50% compared to steady-state walking. Overall, the CoP-mechanism was the main contributor to the total CoM acceleration. The use of the counter-rotation mechanism may be limited because angular accelerations ultimately need to be reversed and because of interference with other task constraints, such as preventing interference with the gait pattern.

## List of symbols and abbreviations

|  |  |
| --- | --- |
| CoM | center of mass |
| BoS | base of support |
| ML | mediolateral |
| CoP | center of pressure |
| $m$ | body mass |
| CoM<sub>vertical</sub> | vertical position of the CoM |
| CoM<sub>ML</sub> | ML position of the CoM |
| $t$ | time |
| $\ddot{\text{CoM}}_{\text{ML}}$ | double derivative of CoM<sub>ML</sub> |
| $g$ | gravitational acceleration |
| CoP<sub>ML</sub> | ML position of the CoP |
| $\dot{H}_{\text{fr}}$ | change in total body angular momentum in the frontal plane |
| $F_z$ | vertical ground reaction force |

## Acknowledgments

The authors are thankful for all participants, Leon Schutte for the technical support and Moira van Leeuwen and Marijne Nieuwelink for assistance during the data collection.

## Data availability

The data that support the findings of this study are available on request from the corresponding author, [SMB].

